# Distinct Modes of Functional Connectivity induced by Movie-Watching

**DOI:** 10.1101/286484

**Authors:** Murat Demirtaş, Adrian Ponce-Alvarez, Matthieu Gilson, Patric Hagmann, Dante Mantini, Viviana Betti, Gian Luca Romani, Karl Friston, Maurizio Corbetta, Gustavo Deco

## Abstract

A fundamental question in systems neuroscience is how endogenous neuronal activity self-organizes during particular brain states. Recent neuroimaging studies have revealed systematic relationships between resting-state and task-induced functional connectivity (FC). In particular, continuous task studies, such as movie watching, speak to alterations in coupling among cortical regions and enhanced fluctuations in FC compared to resting-state. This suggests that FC may reflect systematic and large-scale reorganization of functionally integrated responses while subjects are watching movies. In this study, we characterized fluctuations in FC during resting-state and movie-watching conditions. We found that the FC patterns induced systematically by movie-watching can be explained with a single principal component. These condition-specific FC fluctuations overlapped with inter-subject synchronization patterns in occipital and temporal brain regions. However, unlike inter-subject synchronization, condition-specific FC patterns contained increased correlations within frontal brain regions and reduced correlations between frontal-parietal brain regions. We investigated the condition-specific functional variations as a shorter time scale, using time-resolved FC. The time-resolved FC showed condition-specificity over time, notably when subjects were also watching the same and different movie scenes. To explain the self-organisation of whole-brain FC through the alterations in local dynamics, we used a large-scale computational model. We found that the condition-specific reorganization of FC could be explained by local changes that engendered changes in FC among higher-order association regions, mainly in frontal parietal cortices.

## Introduction

The neural correlates of information processing at a local scale have been widely studied. However, the integration of information at the whole-brain level may also be crucial for understanding brain function (Baars, 1993; Tononi, 2004). Advances in neuroimaging techniques such as functional magnetic resonance imaging (fMRI) now allow us to ask how the brain regulates information flow in large-scale cortical networks (Deco et al., 2015). For example, several studies suggest that neuronal synchronization mediates communication in large-scale cortical networks during task performance (Brovelli et al., 2004; Gross et al., 2004; Siegel et al., 2008) and the resting state (Brookes et al., 2011; Hipp et al., 2012).

Resting state functional connectivity (rs-FC) is a widely-used technique to characterize large-scale organization of brain activity, based on the temporal correlations between blood oxygen level-dependent (BOLD) signals (Biswal et al., 1995). Rs-FC patterns have been shown to provide ‘fingerprints’ for the functional brain organization during the resting-state (Finn et al., 2015; Smith, 2016) and task induced responses (Tavor et al., 2016). Recent studies suggest a strong relationship between the FC during resting state and task performance (Betti et al., 2013; Cole et al., 2016, 2014; Rosenberg et al., 2015). In particular, continuous task paradigms such as viewing natural scenes (i.e. movie watching) are of particular interest due to their ecological validity (Betti et al., 2013). Several studies have found that FC is more reliable and promotes the detection of individual differences while subjects are watching movies (Kim et al., 2017; Vanderwal et al., 2017, 2015). Moreover, a systematic reorganization of the cortical interactions - with changes in functional network assignments - has been demonstrated during movie-watching (Kim et al., 2017; Wolf et al., 2010). Therefore, the condition-specific changes and enhanced reliability of FC may be induced by the task-dependent engagement of specific brain regions (Hasson, 2004; Hasson et al., 2010) and/or large-scale functional reorganization (Kim et al., 2017; Simony et al., 2016; Wolf et al., 2010). On the basis of these studies, we hypothesized that the intrinsic reorganization of FC during movie-watching could be quantified and modelled in terms of systematic fluctuations in connectivity patterns.

To study the reorganization of FC, we analysed the variations in grand-average (over time) and time-resolved FC during rest and movie-watching. We characterized the variations in FC patterns across subjects using principal component analysis (PCA). PCA and associated techniques have been used to characterize resting-state fluctuations (Carbonell et al., 2011), whole-brain connectivity dynamics (Allen et al., 2012) and disease-related rs-FC states (Craddock et al., 2009). Based on the projections of individual subject scores on the principal components, we identified FC-states specific to the movie-watching condition. We then compared these condition-specific FC patterns with inter-subject synchronization (Kim et al., 2017; Simony et al., 2016).

One question related to the task-dependent reorganization of FC is whether alterations in grand-average FC (over the whole session) reflect a continuous (temporally stable) functional state or the emergence of functional modes fluctuating over time (Gonzalez-Castillo et al., 2015). To answer this question, we extended our analysis beyond grand-average FC states and investigated the temporal fluctuations in FC states based on the dynamics of phase-coupling among brain regions.

Finally, we used whole-brain computational modelling to test whether the reorganization of FC can be explained by the fluctuations in local connectivity. In other words, we adopted a mechanistic approach to task-dependent FC using a large-scale, biophysically plausible modelling framework. In brief, we constrained long-range interactions between brain regions using diffusion weight imaging-derived (DWI) structural connectivity, and estimated the fluctuations in local connectivity - of each brain region - during movie-watching that best explained the observed FC.

## Results

To characterize fluctuations in functional connectivity (FC), first we established the relationship between the FC patterns during resting-state and movie-watching conditions. The grand average FC over the resting-state and movie-watching sessions exhibited similar patterns (*r=0.8*) **(Figure 1A)**. The similarity among the FC of individual subjects was substantially higher under the same condition (*resting-state r=0.46 ± 0.06; movie r=0.49 ± 0.06*) than across conditions (*r=0.40 ± 0.07*). These results confirm previous findings that showed similar grand average FC patterns during resting-state and movie-watching (Betti et al., 2013; Cole et al., 2014).

**Figure 1.**
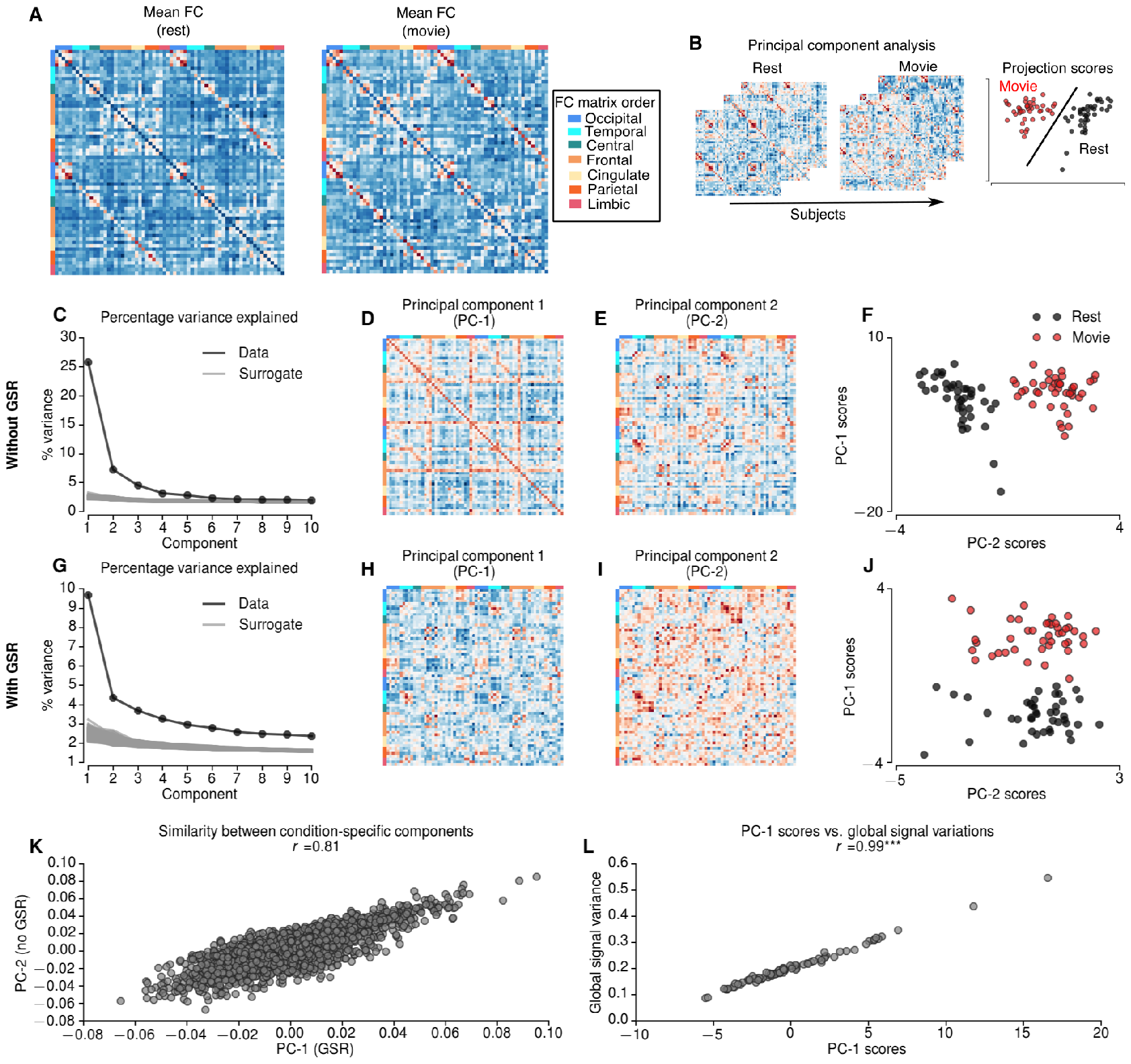
**A** Mean functional connectivity (FC) during resting-state and movie-watching conditions. **B** Schematic describing principal component analysis (PCA) over FCs of 2 resting-state and 2 movie-watching condition concatenated across 21 subjects. **C-F** PCA results without global signal regression (GSR). Explained variance by each PC (black) and random surrogates (gray) without GSR **(C)**. Compared to 1000 random surrogates the dimensionality of FCs without GSR was 13. The first PC **(D)** explains 25.8% of the variation, whereas second PC **(E)** explains 7.2% of the variation. The projections of first two PCs shows that the second component is specific to movie runs **(F)**. The first PC of the FCs without GSR reflects global signal standard deviation **(L)**. **G-J** PCA results with global signal regression (GSR). **G** Explained variance by each PC (black) and random surrogates (gray) with GSR. Compared to random surrogates the dimensionality of FCs with GSR is 22. The first PC, which is specific to movie runs explains 9.69% of the variation **(J)**. **K** The similarity between condition-specific components with and without GSR. *** indicates p<0.0001.

To quantify the variability in FC across subjects during resting-state and movie-watching conditions, we performed principal component analysis (PCA) over subjects **(Figure 1B)**. PCA decomposes high-dimensional data features into orthogonal axes (principal components) that explain the most variance. The projections provide a score for each observation (i.e., subject/run) along the principal components. We concatenated vectorised matrices from 21 subjects, during 2 separate runs of resting state and movie-watching conditions, and then employed PCA. Then, we examined the scores (i.e. expression of principal components by individual subjects during rest movie-watching).

### Distinct modes of variation in functional connectivity during movie-watching

The first principal component (PC-1) - explaining 25.8% of the variance **(Figure 1C)** - reflected a FC pattern that was conserved over runs. The scores of PC-1 were significantly correlated with the global variance of each fMRI run (*r=0.99, p<0.0001, dof=83*) **(Figure 1L)**. This result suggests that the principal mode of variation in FC reflects variations in global signal. The second principal component (PC-2) **(Figure 1E)** - explaining 7.2% of the variance - clearly distinguished the movie-watching condition from resting-state (i.e. given the threshold=0, the PC-2 scores perfectly separates resting-state and movie-watching conditions. We will refer to this component as a condition-specific PC **(Figure 1F)**. This result suggests that the condition-specific variations in FC can be explained along a single mode of variation (PC-2), which is orthogonal to the global-signal related mode (PC-1).

We repeated PCA for 1000 surrogate FCs across subjects to define the components explaining a significant proportion of variance (**see Materials and Methods**). The variance explained by the first 13 components was greater than the variance explained by surrogate FCs; suggesting that the first 2 PCs explain a significant amount of variation. The remaining components did not show any specificity regarding the movie-watching condition and were not analysed further.

To test the consistency of the condition-specific PCs across runs, we repeated the PCA for each run separately and quantified the similarities between PCs across runs. For each run, we identified condition-specific PCs that were highly consistent across runs (*r=0.83 for PC-2 scores*) **(Supplementary Figure 1)**. Furthermore, the similarities between PCs and scores were higher for condition-specific components than global signal-related components (*r=0.75 for PC-1 scores*) **(Supplementary Figure 1)**. These results suggested that the condition-specific PC and associated scores (i.e., expression in individual subjects) were conserved across runs, which suggest a link between condition-specific and individual variations in FC.

### Contribution of potential non-neuronal confounds

Previous studies have shown differential subject movements and increased arousal while watching natural scenes (Siegel et al., 2016; Vanderwal et al., 2015). Therefore, the condition-specific PC may reflect the contributions from movement or arousal artefacts. To address the role of head motion, we calculated the correlation between mean frame-wise displacement and principal component scores. The first PC scores, reflecting global signal variations, were significantly correlated with head motion (*Spearman rank r=0.37, p<0.001, dof=83*). We found no significant correlation between head motion and condition-specific PC scores (*Spearman rank r=0.03, p=0.75, dof=83*).

To preclude other artefactual contributions, we repeated the analyses after regressing out the global signal **(Figure 1G-J)**. After global signal regression (GSR) the first principal component (PC-1) explained 9.69% of the variance and reflected condition-specific variations in FC **(Figure 1J)**. Similarly, no significant correlation was observed between head motion and condition-specific PC scores after GSR (*Spearman rank r=0.02, p=0.85, dof=83*). Crucially, the condition-specific components were similar with and without GSR (*r=0.81*) **(Figure 1K)**. This analysis suggests that condition-specific variations in FC are not associated with head motion and that they are robust to global signal regression.

### Relationship between condition-specific FC variations and inter-subject synchronization

The condition-specific variations in FC may reflect time-locked fluctuations during movie-watching condition as reported in previous studies (Kim et al., 2017; Simony et al., 2016). We characterized these time-locked FC patterns (during the movie-watching condition) using inter-subject synchronization FC (ISS-FC) **(Figure 2A)**. In brief, ISS-FC removes the contribution of endogenous activity by evaluating the FC between two regions from different subjects (Kim et al., 2017; Simony et al., 2016). For each run, the subjects were randomly assigned into 2 non-overlapping groups. The FCs were then evaluated as the correlations between pairs of regions across the average BOLD time-series of distinct set of subjects. Since the subjects were exposed to the same stimuli only during movie watching, ISS-FC exhibited high-magnitude correlations in the movie-watching but not in the resting-state condition (Kim et al., 2017)**(Supplementary Figure 2)**.

**Figure 2.**
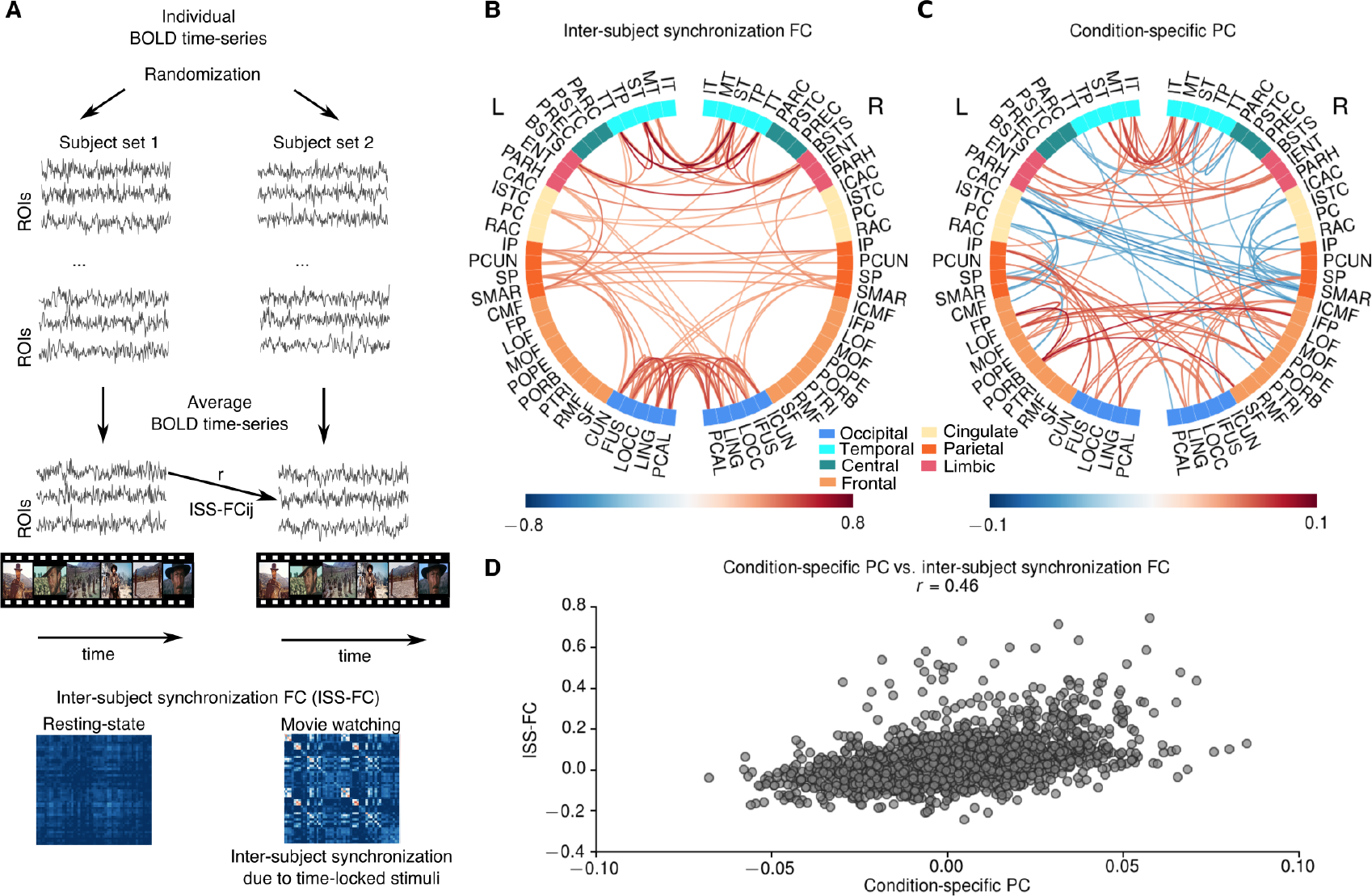
Comparison of condition-specific PC and inter-subject synchronization FC. **A** Schematic illustrating the computation of inter-subject synchronization FC (ISS-FC). The subjects were randomly separated into 2 groups. Then the average BOLD time-series were calculated for each group. ISS-FCs were computed as the correlation between BOLD time-series across groups for each pair of regions. **B** The largest 100 connections in ISS-FC during movie-watching condition. The most prominent correlations were observed among occipital and temporal brain regions, and between occipital and parietal brain regions. **C** The largest 100 connections in condition-specific PC. Condition-specific PC also shows increased connectivity among occipital and temporal brain regions, and between occipital and parietal brain regions; and the overall connectivity pattern in ISS-FC and condition-specific PC was highly similar **(D)**. However, the condition-specific PC also exhibited increased connectivity among frontal brain regions and suppressed connectivity between cingulate and parietal regions **(C)**.

ISS-FC during movie watching showed the highest values within occipital and temporal regions; suggesting that synchronization is due to time-locked visual and auditory events **(Figure 2B)**. In addition, ISS-FC showed high synchronization between occipital/temporal and parietal brain regions, such as inferior and superior parietal cortex **(Figure 2B)**. The pattern of the condition-specific PC was similar to the ISS-FC (*r=0.46*) **(Figure 2C-D)**. As in the ISS-FC, the condition-specific PC exhibited higher values within occipital and temporal, and between occipital/temporal and parietal brain regions **(Figure 2C)**. However, the condition-specific PC differed from the ISS-FC in various aspects: First, the condition-specific PC exhibited more pronounced connectivity changes in fusiform and lingual gyri, and inferior temporal compared to the ISS-FC. Second, the condition-specific PC comprised enhanced intra- and inter-hemispheric connectivity between frontal brain regions (particularly lateral and medial orbital frontal cortex, pars orbitalis and frontal pole), which were not observed in the ISS-FC. Third, the condition-specific PC exhibited strong negative values (reduced connectivity) particularly across frontal and parietal regions. These attenuated values involved FC between caudal anterior/posterior cingulate and supramarginal gyrus, superior/inferior parietal, and caudal middle-frontal cortex. These results suggest that although the condition-specific PC overlaps with the ISS-FC, it highlights a distinct functional reorganization, expressed predominantly in higher-order association regions.

### Condition-specific FC trajectories in time-resolved FC

The grand average FC approach cannot disambiguate between a temporally stable mode of FC and fluctuations in FC (i.e., a succession of distinct FC patterns). To address this issue, we analysed time-resolved fluctuations in FC (also known as dynamic FC). Here, we tested the hypothesis that FC continuously reorganizes during movie-watching. We constructed time-resolved FC based on the fluctuations in phase-locking values (PLVs) between brain regions (see Materials and Methods). The advantage of this approach is that it eliminates the dependency on a particular window and step size, as in sliding-window analysis. Instead, it requires one to specify a narrowband range to calculate PLVs. Here, we chose *0.04-0.07Hz*, which does not overlap with the frequency ranges of low-frequency drift and high-frequency noise (Glerean et al., 2012). First, we band-pass filtered the BOLD time-series and employed Hilbert transform. We then calculated the PLVs at each time point using the instantaneous phases of each brain region **(Figure 3A)**.

**Figure 3.**
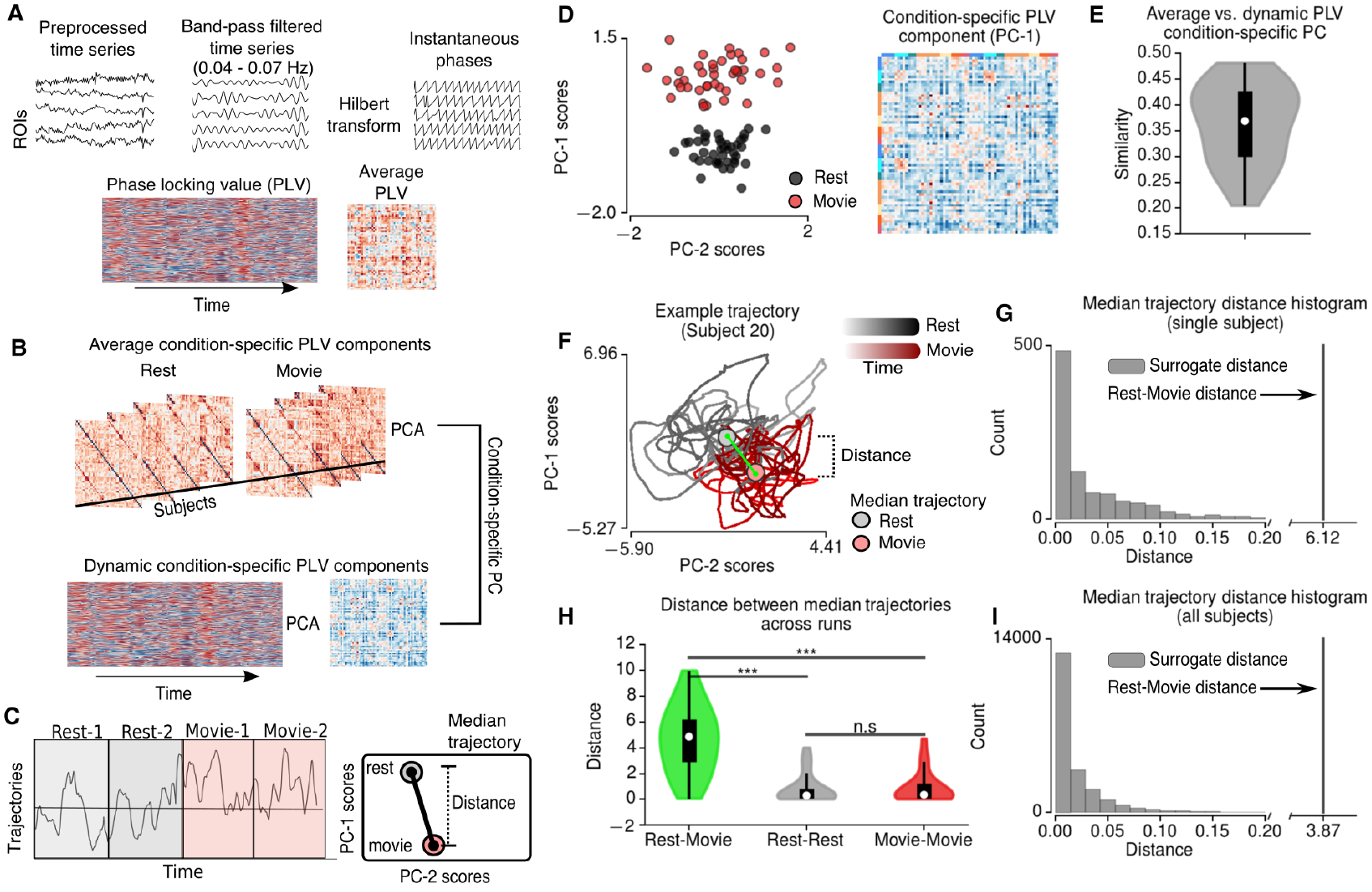
Time-resolved FC based on phase locking values (PLVs). **A** Schematic describing the calculation of PLVs. Preprocessed BOLD time-series were narrow-band filtered in 0.04-0.07Hz range and the resulting signals were Hilbert transformed. Phase-locking values were calculated based on the difference between instantaneous phases across brain regions. **B** Schematic describing principal component analysis (PCA) performed on average PLVs across subjects (top) and dynamics of PLVs across time for each subject (bottom). A condition-specific component was identified based on the maximum similarity between dynamic PLV components and average condition-specific PLV component **(D)**. The average and dynamic condition-specific components were very similar across subjects **(E)**. Based on the trajectories of condition-specific PLV components, the distance between the median trajectories of resting-state and movie-watching conditions were calculated **(C)**. **F** Example trajectory for single subject. **G** The distance between the median trajectories of resting-state and movie-watching conditions compared to the histogram of the distances for 1000 randomly split trajectories. **H** The median trajectory distances between resting-state and movie-watching conditions, between 2 resting-state runs and between 2 movie-watching runs. The distance between conditions was significantly higher than the distance between runs (permutation t-test, 10000 permutations). *** indicates p<0.0001, n.s. indicates p>0.05.

To establish the link between the time-resolved FC analyses and Pearson correlation-derived FC, we calculated the grand average PLVs over time, and performed PCA over subjects. This analysis showed that the principal components based on PLVs also exhibit condition specificity **(Figure 3D)**. Furthermore, condition-specific PC of PLVs was similar to those derived from Pearson correlation-derived FC (*r=0.88*). Therefore, the condition specific FC patterns for PLVs were aligned with those based on the Pearson correlation-derived grand-average FC.

For each subject we performed PCA on PLVs over time **(Figure 3B)**. We identified condition-specific component for each subject as the one (i.e. PC-1 or PC-2) exhibiting the highest correlation with the grand-average condition-specific component (**Figure 3E)**. For the majority of the subjects, the trajectories (i.e. the PC scores) of the condition-specific components reflected a clear distinction between conditions **(Supplementary Figure 3)**. We quantified this condition-specificity for each individual subject by comparing the median trajectories (i.e. median PC scores) during the resting-state and the movie-watching conditions **(Figure 3C)**. We then calculated the distance (i.e. squared difference) between the median trajectories of rest/movie conditions **(Figure 3F)**. The distance between rest/movie median trajectories were compared to the distance between 1000 randomly grouped trajectories **(Figure 3G)**. 20 out of 21 subjects showed a significantly larger distance between rest/movie trajectories than any other randomly grouped trajectories (*p<0.001*) **(Figure 3I)**. Since the trajectories of the condition-specific PCs are time-dependent, we assessed the significance of the median trajectory distances between runs/conditions across subjects. We found that the distance across conditions (i.e. movie/rest conditions) were significantly larger than the distance across runs (i.e. rest/rest and movie/movie runs) (*p<0.0001, permutation t-test, 10000 permutations*) **(Figure 3H)**. We found no significant difference between the distance across runs for resting state and movie conditions (*p=0.82, permutation t-test, 10000 permutations*) **(Figure 3H)**. These results speak to the emergence of a conserved FC pattern during movie-watching condition on a short timescale.

### Condition-specific FC patterns within and across runs

To study the role of time-locked events on PLV dynamics during movie-watching (analogous to inter-subject synchronization), we calculated the similarity between instantaneous PLVs across conditions and runs. In brief, for each time point, we calculated the similarity between the PLVs of a single subject (k) and the average PLVs across the rest of the subjects (n≠k). The average PLVs were calculated to test the PLV similarity in 3 different cases: Across conditions (e.g. if subject k is at resting state run 1, the average PLVs were calculated for movie-watching run 1), across runs (e.g. if subject k is at resting state run 1, the average PLVs were calculated for resting state run 2) and within runs (e.g. if subject k is at resting state run 1, the average PLVs were calculated for resting state run 1) **(Figure 4A)**.

**Figure 4.**
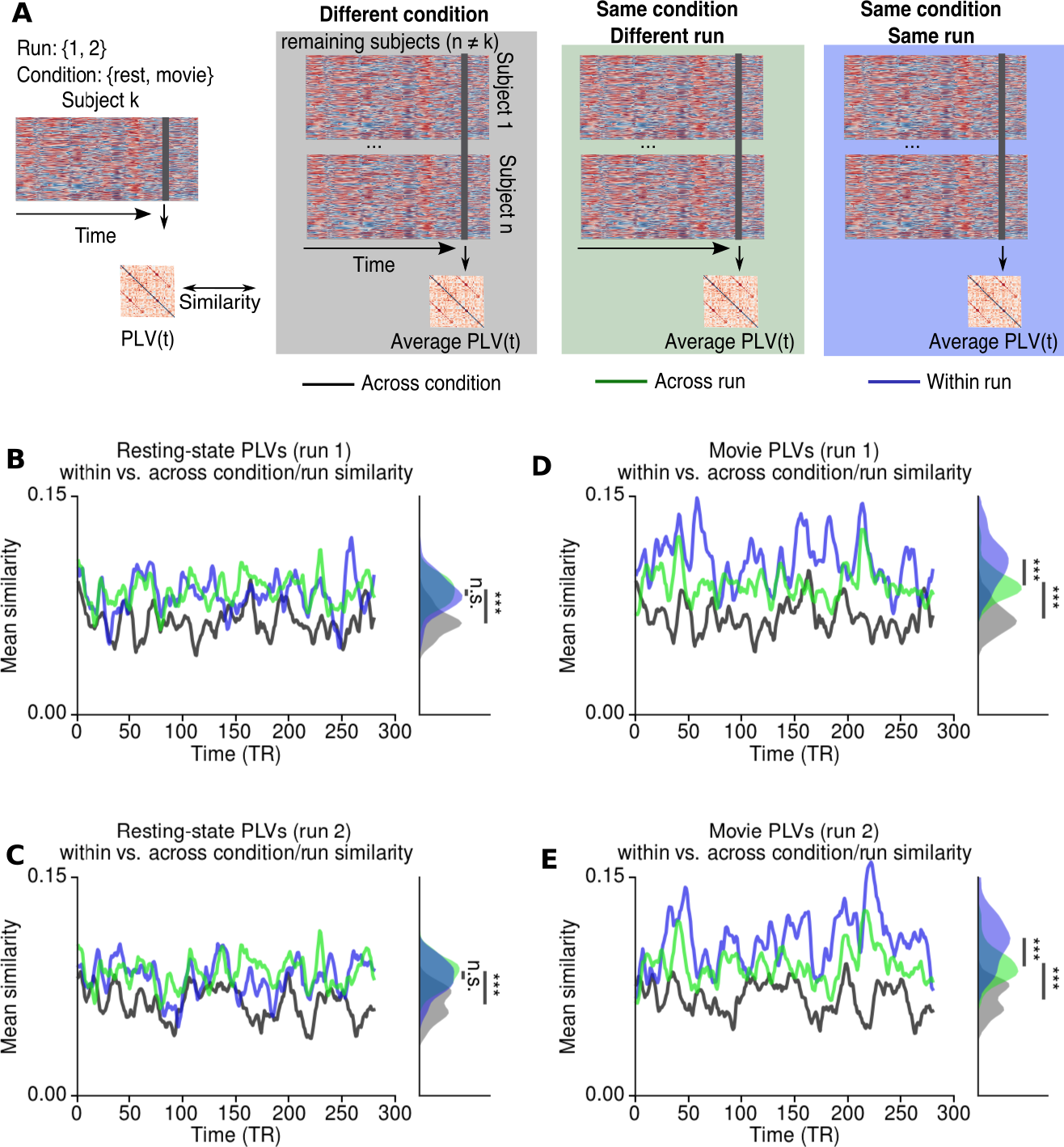
Time-resolved similarity between PLVs across conditions and runs. **A** Schematic describing the procedure. For each subject the PLVs at each time point was compared to the average PLVs across the remaining subjects at the same time point. Black/gray lines/shades indicate that the average PLVs were calculated for different condition (i.e. if subject k is at rest, average PLVs were calculated for movie-watching). Green lines/shades indicate that the PLVs were calculated for the same condition but different run (i.e. if subject k is at rest in run 1, average PLVs were calculated for the resting-state run 2). Blue lines/shades indicate that the PLVs were calculated for the same condition and the same run (i.e. if subject k is at rest in run 1, average PLVs were calculated for the resting-state run 1). **B-C** During resting-state the similarity between PLVs were significantly lower across conditions (i.e. rest vs. movie), but there was no significant difference between the similarities across runs. **D-E** During movie-watching, the similarity between PLVs was significantly lower across conditions. However, the similarity between PLVs was significantly higher within runs compared to across runs. The histograms illustrates the distributions of similarity measures over time, whereas *** indicates the p<0.0001 assessed by permutation t-test across subjects. n.s. indicates p>0.05.

Both for resting-state and movie-watching conditions, the similarity across runs was significantly higher than the similarity across conditions (*p<0.0001 for both runs, permutation t-test, 10000 permutation*) **(Figure 4B-E)**, confirming the continuous functional reorganization during movie-watching condition. Furthermore, this result shows that during movie-watching condition, the similarity between instantaneous PLVs was higher, even when the subjects were viewing different scenes. For resting state runs, the average similarity between instantaneous PLVs did not show any significant difference across runs (*p=0.54 for run 1, p=0.34 for run 2, permutation t-test, 10000 permutations*) **(Figure 4B,C)**. In contrast, the average similarity between instantaneous PLVs were significantly higher for the same movie runs than across runs (*p<0.0001 for both runs, permutation t-test, 10000 permutation, p<0.0001*) **(Figure 4D,E)**. These results indicate that the PLV dynamics during movie-watching reflects both the effects of time-locked events and a continuous functional reorganization.

### Large-scale computational modelling of the regional dynamics underlying movie-watching FC

Both the grand average and time-resolved FC analyses suggested a functional reorganization during movie-watching. Based on these results, we hypothesized that the variations in regional dynamics could explain the functional reorganization. We used Hopf normal model to characterize the BOLD activity of each region (Deco et al., 2017). The regions were coupled to each other via DWI-derived structural connectivity scaled by a global coupling parameter **(Figure 5A)**. The dynamics of each region were governed by a local bifurcation parameter (a). The local bifurcation parameters (a) reflect whether an individual region is in a noise-driven regime (a < 0), oscillatory regime (a > 0), or alternates between the two regimes (a ~ 0) **(Figure 5A)**. We estimated the global coupling and local bifurcation parameters of each subject/run by maximizing the similarity (i.e. Pearson correlation) between empirical and model FCs using gradient-descent. We found no significant difference between the model fits for resting-state (*r=0.518 ± 0.057*) and movie-watching conditions (*r=0.497 ± 0.045*) (*p=0.146, permutation t-test*, 10000 permutations). To characterize the overall topography underlying each condition, first we estimated the optimal global coupling parameter (g) and optimal bifurcation parameters (a) for resting state and movie watching condition. At rest, the average bifurcation parameter estimates were low in parietal and temporal regions, whereas they were higher in occipital and frontal regions **(Figure 5B)**. For movie condition, the bifurcation parameters were elevated in parietal and temporal regions and decreased in anterior cingulate, lateral prefrontal cortices and in supramarginal gyrus **(Figure 5C)**. There was no difference between the mean optimal bifurcation parameters of rest and movie conditions **(Figure 5D)**.

**Figure 5.**
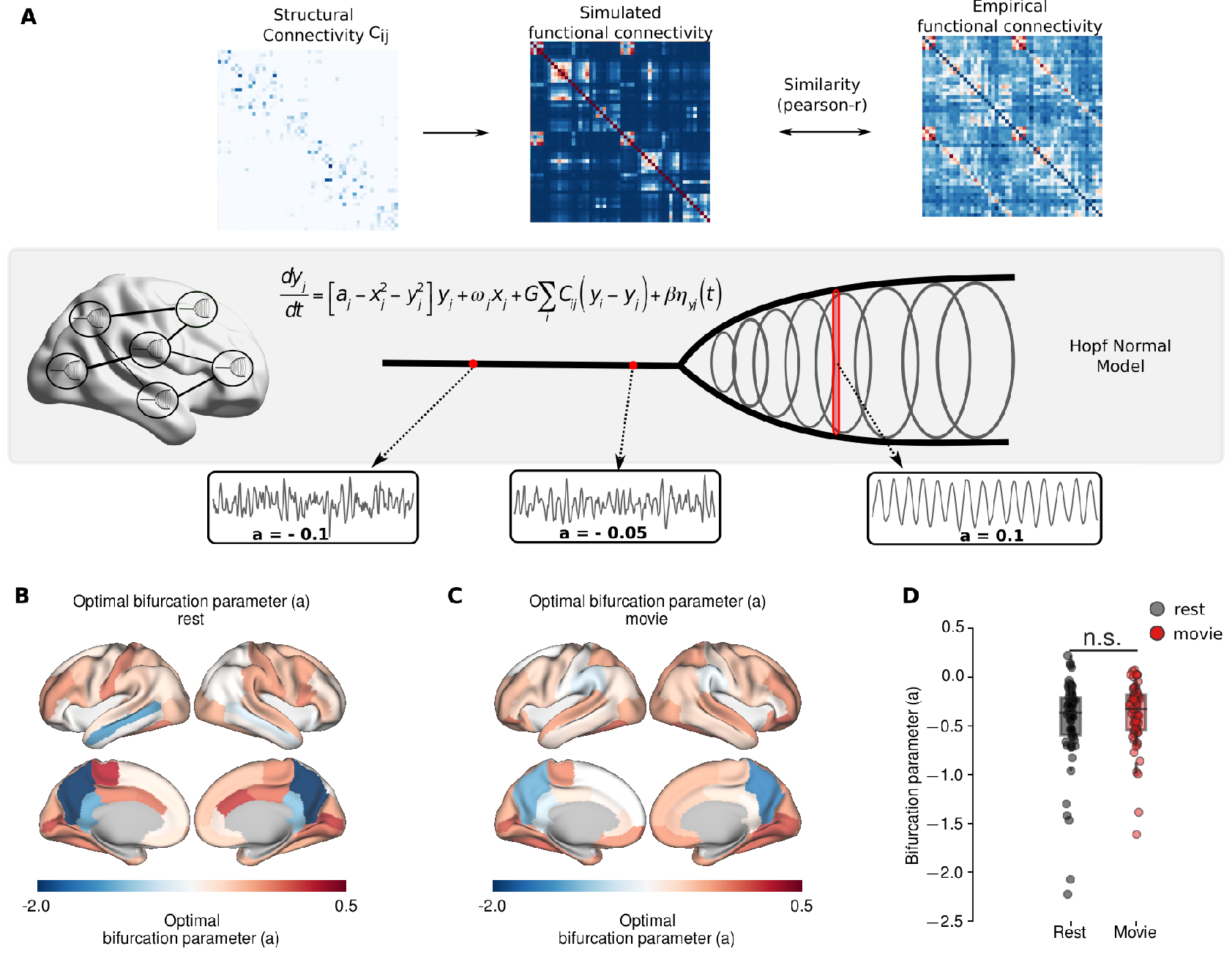
Large-scale computational modelling. **A** The schematic of the modelling framework. The BOLD activity of each region was described using Hopf normal model, where the local bifurcation parameters (a) mediate the local dynamics. Negative values of bifurcation parameter, a, indicates noise-driven activity, whereas positive values indicate oscillatory activity with increasing amplitude. Brain regions are coupled each other through DWI-derived SC matrix. The optimal model parameters were estimated using gradient descent optimization, which maximizes the similarity between empirical and model FC. **B** Mean optimal bifurcation parameter topography at resting state. **C** Mean optimal bifurcation parameter topography during movie condition. **D** The distributions of the bifurcation parameters during movie condition and resting state. n.s. indicate p>0.05.

To quantify the difference between conditions, we compared the optimal global coupling and bifurcation parameters of the resting-state and the movie-watching conditions **(Figure 6A)**. We found no significant difference in global coupling parameters between rest and movie conditions (*p=0.719, permutation t-test*, 10000 permutations) **(Figure 6B)**. In the movie condition, the local bifurcation parameters were significantly decreased - towards negative values - in bilateral caudal anterior cingulate, right supramarginal gyrus, and left postcentral cortex **(Figure 6D)**. In contrast, the bifurcation parameters were significantly increased in bilateral orbital frontal and lateral orbital frontal cortices, left medial temporal cortex, right frontal pole, middle rostral frontal and superior parietal cortex cortices **(Figure 6D)**. These changes in higher-order association regions are consistent with the patterns observed in condition-specific PC.

**Figure 6.**
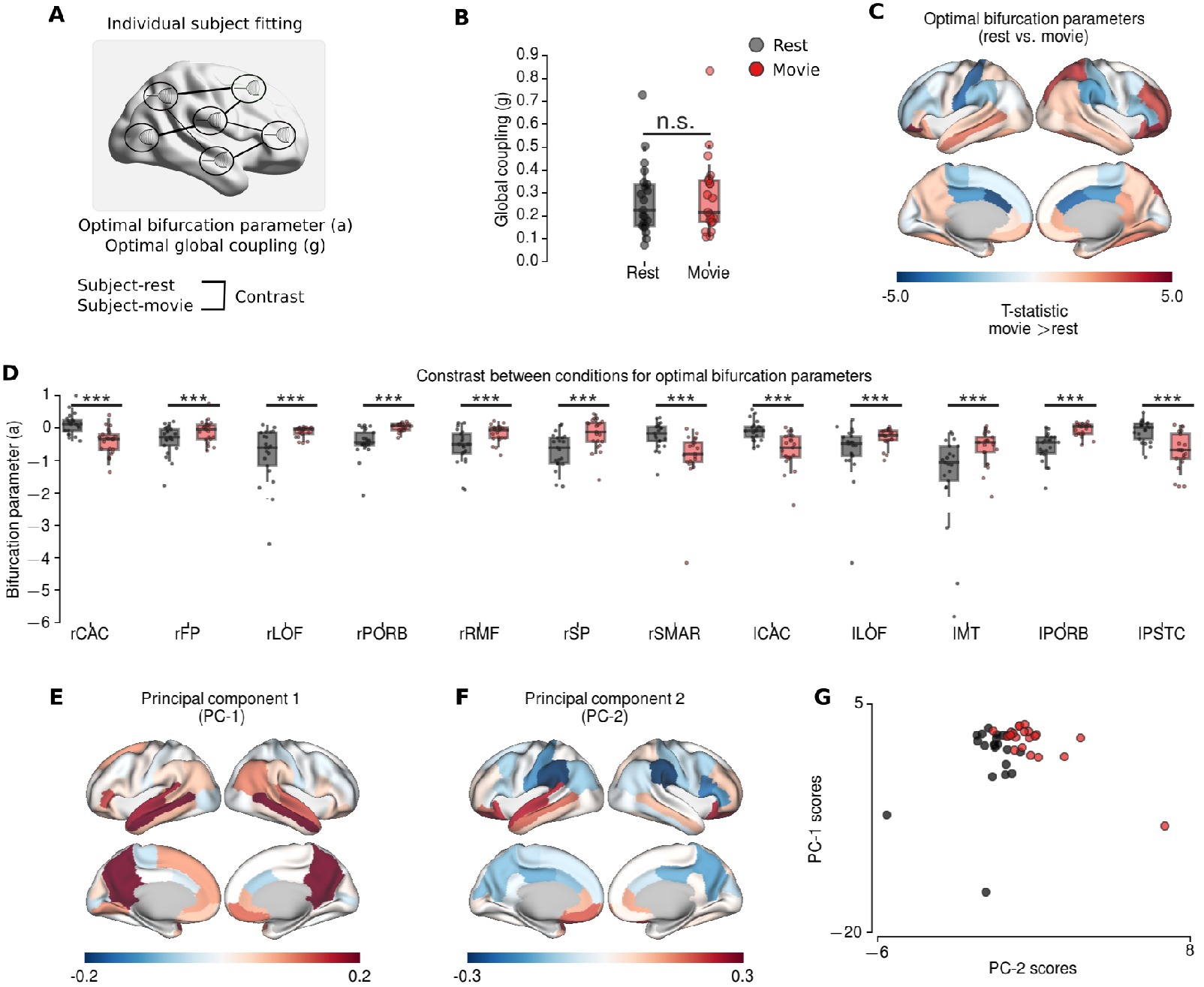
Modelling results for individual subject fitting. **A** The schematic of individual subject fitting. **B** The group differences for global coupling parameters did not show significant difference. **C-D** The group differences between optimal bifurcation parameters at rest (black) and during movie condition (red) (permutation t-test, 10000 permutations). **C** The topography of the group differences (T-statistics; hot colours indicate larger values during movie condition). **D** Boxplots of the regions showing significantly difference after FDR correction (*p<0.01*). **E-G** Principal component analysis applied to optimal bifurcation parameters in the model. **E** The topography of the first principal component. **F** The topography of the second principal component. PC-1 has higher values in precuneus, posterior cingulate, medial temporal and frontal regions, exhibiting typical pattern associated to default mode network. PC-2 exhibit increased values in frontal and temporal regions, and decreased values particularly in supramarginal gyrus consistent with the contrast between conditions. **G** The projections of the principal components on rest and movie conditions. *** indicates p<0.01, n.s. indicates p>0.05.

Finally, we repeated the PCA on the bifurcation parameter estimates across subjects and conditions **(Figure 6E-G)**. The scores of the first principal component (PC-1) - explaining 41.77% of variance - and the second principal component (PC-2) - explaining 10.25% of variance - were both significantly correlated with the scores of the empirically observed condition-specific PC (*PC-1 Spearman rank r=0.44, p=0.004, dof=41; PC-2 Spearman rank r=0.63, p<0.0001, dof=41; PC-1+PC-2 Spearman rank r=0.73, p<0.0001, dof=41*). The first principal component (PC-1) exhibited a strong positive peak in precuneus and isthmus of cingulate; with slightly higher values in medial frontal and temporal regions, which is very similar to default mode network (DMN) topography **(Figure 6E)**. The second principal component (PC-2) had higher values in temporal and frontal regions as observed in the contrast between conditions **(Figure 6F)**. Furthermore, the scores of the first and second principal components were negatively correlated only in movie-watching condition (*Spearman rank r=-0.496, p=0.02, dof=20*) **(Figure 6G)**. These results suggest that the changes in local connectivity during the movie-condition engender multiple modes of variation, which reflect condition-specific and DMN-like topographies.

## Discussion

In this paper, we investigated the reorganization of functional connectivity (FC) during movie-watching condition. We showed that during movie-watching FC patterns vary along a single mode of variation (i.e. a condition-specific pattern of connectivity that captures the variations across subjects), which emerges as a continuous functional state over time.

We used principal component analysis (PCA) to characterize the variations in FC across individuals and conditions (i.e. resting-state vs. movie-watching). We found that the principal component (PC-1) reflected the variations in global signal, whereas the second principal component (PC-2) reflected the distinction between resting-state and movie-watching conditions. We investigated the patterns of the condition-specific component in the context of inter-subject synchronization FC (ISS-FC) (Kim et al., 2017; Simony et al., 2016). The connectivity patterns of the condition-specific component were similar to the ISS-FC. Both characterizations of FC highlighted intra- and inter-hemispheric connectivity within occipital and temporal regions as well as their connections with parietal regions. These results suggest that the enhanced communication between regions related to audiovisual processing and attention are primarily driven by the time-locked events during movie-watching (Hasson, 2004; Hasson et al., 2010).

However, unlike ISS-FC, condition-specific component showed enhanced connectivity within frontal brain regions and reduced connectivity between frontal-parietal brain regions and cingulate (e.g. supramarginal gyrus, superior/inferior parietal cortex, caudal middle frontal cortex vs. anterior and posterior cingulate cortex). These results are consistent with previous studies of functional reorganization during movie-watching (Kim et al., 2017; Simony et al., 2016; Wolf et al., 2010). We argue that - during movie-watching - reorganization of FC with the primary sensory regions is mainly driven by extrinsic factors such as sensory stimulation, whereas the higher-order association regions exhibit a self-organisation of endogenous activity.

The existence of a condition-specific component in grand-average FC may not be sufficient to draw conclusions about the functional reorganization during movie-watching. Therefore, we asked how the condition-specific PC topography relates to the time-resolved FC. We used the Hilbert transform of narrowband filtered BOLD time-series, and characterized time-resolved FC based on phase-locking values over time. We found condition-specific components on grand average PLVs over subjects as well as individual PLVs over time. The trajectories of the condition-specific PLV components suggested that this component might appear as a stable state during movie-watching. We endorsed this conclusion by analysing the similarity between instantaneous PLVs and average PLVs (over subjects), under different conditions/runs. The similarity was significantly lower when the subjects were scanned under different conditions (i.e. rest vs. movie) than they were under same condition (i.e. rest vs. rest and movie vs. movie). Furthermore, only during movie-watching, did we find that PLV similarity was higher for subjects in the same run (i.e. run 1 vs. run 1) than subjects in the different runs (i.e. run 1 vs. run 2). Overall, these results suggested that whole-brain FC (in the time-scale of BOLD signals) is continuously reconfigured on a short time scale. We speculate that the functional reorganization in higher-order association regions may reflect the adaptation of the brain’s intrinsic architecture to mediate large-scale information flow during movie-watching.

Previous studies have reported decreased head movements and higher arousal while movie-watching (Siegel et al., 2016; Vanderwal et al., 2015). Therefore, the emergence of a condition-specific FC component could also reflect systematic artefacts. In this study, we found no significant differences between mean frame-wise displacements (head motion) of the subjects across conditions. However, we observed that head motion was significantly altered while watching movie (i.e. during movie-watching condition some subjects moved less, whereas others move more). The scores of the condition-specific PC were not correlated with the mean frame-wise displacements or the PC scores associated with head motion. However, both measures were significantly correlated with the PC scores reflecting global signal variations. To rule of the possibility of other confounds, we repeated the analysis and identified similar condition-specific component after performing global signal regression (GSR), which was replicated across runs. Apart from the head motion and global signal analyses, the contribution of non-neuronal confounds is unlikely, given the results: First, the variations in occipital and temporal regions in condition-specific component substantially overlaps with inter-subject synchronization (no changes were observed in somatomotor brain regions), which relies on the covariation between brain regions averaged over different subjects. Although the sensory-motor brain regions are known to be more susceptible to non-neuronal confounds (Bijsterbosch et al., 2017; Power et al., 2017), these results are more likely explained by common audiovisual stimulation than synchronization of head motion or respiration across subjects. Second, the condition-specific components were very similar across runs, and narrowband filtered data (0.04-0.07Hz). Known artefactual sources such as low-frequency drift, cardiac and respiratory variations are often associated with lower or higher frequencies (Glerean et al., 2012). Therefore, substantial variations in the condition-specific component would be expected in the narrow-band signals, if they were related to these confounding factors.

Our results suggested a distinct and continuous reorganization of FC during movie-watching. Under the assumption that structural connectivity does not change, one can use whole-brain computational modelling to characterize local variations in neurodynamics during movie-watching. Here, we used Hopf normal model to characterize BOLD signals. The motivation behind using this model was that noise-driven and oscillatory dynamics can be modelled using a single parameter (local bifurcation parameter). When the local bifurcation parameter of a particular region is negative, each region exhibits noise-driven dynamics. For positive bifurcation parameter values, the region exhibits sustained oscillations. Therefore, higher parameters values of a region in the model indicate that the region has larger influence on its connected regions. The model revealed significant decreases in bifurcation parameters particularly in anterior cingulate cortex and in supramarginal gyrus, which suggested an association between decreased bifurcation parameters and the key regions that exhibited suppressed connectivity patterns in the component-specific PC. In contrast, the bifurcation parameters increased in lateral prefrontal cortex, medial temporal cortex and superior parietal regions. These results suggest that endogenous activity in higher-order association regions are altered during movie-watching. Nevertheless, it is important to note that the model describes the BOLD signals in the associated low-frequency narrow-band. Therefore, the results should be interpreted only in relation to low-frequency fluctuations in BOLD signals.

The PCA over model parameters revealed two different modes of variation that were associated with the FC condition-specific variations. Although the second PC was more consistent with the alterations in empirical and model data, the first PC also showed substantial conditional-specificity. Interestingly, the first PC exhibited a pattern typical of default mode network (DMN), which involves the isthmus cingulate, precuneus, medial frontal and temporal cortices. Furthermore, the associated PC scores were negatively correlated across subjects in only the movie-watching condition. Therefore, the model predicts that the interaction between condition-specific and DMN-like activation patterns has a crucial role in the reorganization of FC. However, based on these results, it is not possible to draw conclusions on the causal mechanisms that drive the relationship between DMN and condition-specific networks. The most important caveat is the lack of individual-specific estimates for structural connectivity. Therefore, the emergence of DMN-like component may simply reflect an additional mode of variation that compensates for the lack of variability in individual-specific structural connections. However, the results may also indicate that several regions of DMN (particularly precuneus) having a role in mediating the switch between distinct functional states, which is consistent with previous studies showing that precuneus dynamically binds to distinct functional networks (Utevsky et al., 2014). An alternative explanation may involve the variations of arousal and vigilance levels. This explanation is consistent with a selective neuromodulatory enabling of intrinsic synaptic connections by ascending modulatory neurotransmitter systems (e.g., noradrenaline) (Shine et al., 2018). This is particularly relevant in light of the systematic changes in the local bifurcation parameter that showed regionally-specific and condition-sensitive effects in our modelling analyses. Recent studies showed the relationship between transcriptomic variations and task-related alterations (Shine et al., 2018) as well as microcircuit specialization (Burt et al., in Press) in the human brain. These advances may allow systematic investigation of the mechanisms behind the functional reorganization of the brain.

Finally, several limitations should be noted while interpreting the results in this paper. The most important limitation of this study is the small sample size (21 subjects). Therefore, the results require replication in an independent dataset. In addition, the design of this study did not allow us to compare the results with other conditions (such as a different movie). Although different runs involved different scenes of the same movie, previous studies have found differences in FC regarding the type/familiarity of the movie (i.e. abstractness, social content) (Vanderwal et al., 2015; Wolf et al., 2010). Future studies may investigate the variants of the movie-watching condition, different tasks and/or other continuous experimental paradigms (e.g. reading, social interactions, etc.). Another important limitation of this study is the use of coarse (33 regions per hemisphere), anatomically defined parcellation. Recently developed cortical parcellations offer functional (or multimodal) definitions of cortical areas, which also facilitate better mapping of functional networks. Our coarse parcellation of cortex had advantages particularly for time-resolved FC analysis and whole-brain modelling due to computational efficiency and the implicit reduction in the number of parameters. Techniques such as independent component analysis may provide better characterization of time-dependent states. Such analytical extensions would require longer recording sessions and a better definition of the cortical areas. A limitation regarding the computational modelling is that the model relies on average DWI-derived SC, which may fail to detect interhemispheric connections, individual variations, and is insensitive to directed connections. Previous studies have shown that the changes in directed effective connectivity may also play role in defining the reorganization of FC (Gilson et al., 2017), which may explain lack of significant differences in visual cortex. Effective connectivity - as assessed using dynamic causal modelling studies of the resting state - also point to a modulation of regional excitability by different components of the default mode. For example, previous studies revealed that the influence of the SN (salience network) and DAN (dorsal attention network) on the DMN (default mode network) regions is inhibitory; whereas the DMN exerted an excitatory influence on the SN and DAN regions (Zhou et al., 2018).

Current experimental paradigms are optimal to study task-dependent changes in BOLD signals, but these may not reveal the dynamic organization of whole-brain FC. Unlike other task-evoked experimental approaches, continuous task paradigms offer a contextual environment (e.g. movie-watching), which engage a collection of processes (e.g. audiovisual processing, attention, social cognition…etc.) contextualized by the stimuli. Our findings suggest that continuous task experiments may be useful to study how humans hierarchically reorganize its internal representations to adapt to environmental context (Friston, 2010). Impairments in these adaptation mechanisms may explain the symptoms in various mental disorders such as schizophrenia (Stephan et al., 2016). Future studies with more sophisticated continuous experimental designs may reveal richer dynamical manifestation of functional reorganization such as consolidation of particular functional states in time (i.e. adaptation) and/or emergence of observable transient functional states (i.e. multistability).

## Materials and Methods

### Study design

The fMRI imaging data used in this paper have been described in details elsewhere (Betti et al., 2013; Mantini et al., 2012). Twenty-four right-handed young, healthy volunteers (15 females, 20–31 years old) participated in the study. They were informed about the experimental procedures, which were approved by the Ethics Committee of the Chieti University, and signed a written informed consent. The study included a resting state and a natural vision condition. In the resting state, participants fixated a red target with a diameter of 0.3 visual degrees on a black screen. In the natural-vision condition, subjects watched (and listened) to 30 minutes of the movie “The Good, the Bad and the Ugly” in a window of 24×10.2 visual degrees. Visual stimuli were projected on a translucent screen using an LCD projector, and viewed by the participants through a mirror tilted by 45 degrees. Auditory stimuli were delivered using MR-compatible headphones.

### Data acquisition

Functional imaging was performed with a 3T MR scanner (Achieva; Philips Medical Systems, Best, The Netherlands) at the Institute for Advanced Biomedical Technologies in Chieti, Italy. The functional images were obtained using T2*-weighted echo-planar images (EPI) with BOLD contrast using SENSE imaging. EPIs comprised of 32 axial slices acquired in ascending order and covering the entire brain (32 slices, 230 × 230 in-plane matrix, TR/TE=2000/35, flip angle = 90°, voxel size=2.875×2.875×3.5 mm3). For each subject, 2 and 3 scanning runs of 10 minutes duration were acquired for resting state and natural vision, respectively. Each run included 5 dummy volumes - allowing the MRI signal to reach steady state, and an additional 300 functional volumes that were used for analysis. Eye position was monitored during scanning using a pupil-corneal reflection system at 120 Hz (Iscan, Burlington, MA, USA). A three-dimensional high-resolution T1-weighted image, for anatomical reference, was acquired using an MP-RAGE sequence (TR/TE=8.1/3.7, voxel size=0.938×0.938×1 mm3) at the end of the scanning session.

### Data preprocessing

Data preprocessing was performed using SPM5 (Wellcome Department of Cognitive Neurology, London, UK) running under MATLAB (The Mathworks, Natick, MA). The preprocessing steps involved the following: (1) correction for slice-timing differences (2) correction of head-motion across functional images, (3) coregistration of the anatomical image and the mean functional image, and (4) spatial normalization of all images to a standard stereotaxic space (Montreal Neurological Institute, MNI) with a voxel size of 3×3×3 mm3. Furthermore, the BOLD time series in MNI space were subjected to spatial independent component analysis (ICA) for the identification and removal of artefacts related to blood pulsation, head movement and instrumental spikes (Smith et al., 2010). This BOLD artefact removal procedure was performed by means of the GIFT toolbox (Medical Image Analysis Lab, University of New Mexico). No global signal regression or spatial smoothing was applied to the preprocessed time series. For each recording (subject and run), we extracted the mean BOLD time series from the 66 regions of interest (ROIs) of the brain atlas (Hagmann et al., 2008)**(Supplementary Table 1)**. We avoided regressing out nuisance parameters and using motion scrubbing, because the effects of these procedures on time-resolved FC analyses (phase locking values) could be unpredictable. 2 subjects were excluded due to signal dropout and 1 subject was excluded due to substantial spikes in the time-series.

### Anatomical Connectivity

Anatomical connectivity was estimated from Diffusion Spectrum Imaging (DSI) data collected in five healthy right-handed male participants (Hagmann et al., 2008; Honey et al., 2009). The grey matter was first parcellated into 66 ROIs, using the same low-resolution atlas used for the FC analysis. For each subject, we performed white matter tractography between pairs of cortical areas to estimate a neuroanatomical connectivity matrix. The coupling weights between two brain areas were quantified using the fiber tract density, and were proportional to a normalized number of detected tracts. The structural matrix (SC) was then obtained by averaging the matrices over subjects.

### Principal component analysis

For all subjects and runs (i.e. 21 subjects, 2 resting state and 2 movie runs) the functional connectivity matrices were constructed based on Pearson correlation coefficient between all pairs of ROIs.

The upper triangular parts of FC (i.e. 66(66 − 1)/2 connections) matrices were concatenated across subjects/runs (21×4 subjects/runs) leading to the feature matrix with dimensions 2145 × 84 (number of connections x number of subjects/runs). Then, principal component analysis (PCA) was applied to the resulting feature matrix. To identify the noise components, the analyses were repeated for 1000 surrogate time-series for each subject/run. The properties of the surrogate time-series of each individual subject were preserved in spectral domain (Prichard and Theiler, 1994). The dimensionality of the data was characterized by the fraction of explained variance of the principal components that are larger than those of the surrogates. Since PCA decomposes the data into orthogonal axes with associated projections (i.e. scores) of each individual observation, we characterized the components based on these projections scores. The first PC might reflect the global synchronization levels. To quantify this, we calculated the correlation between the first PC scores and the variance of global signal (i.e. the mean signal across regions). The principal component related to movie-watching condition was characterized as the one exhibiting clear separation between conditions based on the PC scores (i.e. the scores higher than 0 indicated the movie-watching runs, whereas the scores less than 0 indicated the resting-state runs).

To quantify the consistency of principal components, we repeated the analysis using 2 separate runs. For both runs, the feature matrices comprised the concatenated upper triangular FC matrices of 1 resting state run and 1 movie run (i.e. 2145 × 42 matrices). The consistency was quantified as Pearson similarity of the components and their projections across runs (Supplementary Figure 1).

### Non-neuronal confounds

During natural viewing condition the individuals are shown to have restricted movements and increase arousal (Vanderwal et al., 2015). Therefore, the differences in FC can be substantially affected by underlying artefacts. For each subject and run, we quantified head motion by calculating the mean frame-wise displacement (Power et al., 2012). We found no significant differences in mean frame-wise displacement across conditions (*p=0.21, permutation t-test, 10000 permutations*). However, we observed condition-specific changes in motion parameters (i.e. several subjects consistently exhibited higher head movement during movie-runs, whereas other subjects exhibited lower head movement). To test this observation, we first used a regression model for mean frame-wise displacement:

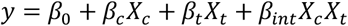

Where mean frame-wise displacement is *y*, *X*_*c*_ is a dummy variable representing condition (resting-state vs. movie-watching), *X*_*t*_ is another variable representing each subject’s tendency to exhibit increased/decreased movement during movie condition. The regression coefficient was not significant for the condition term (*p=0.71*), but the coefficients were significant for tendency and the interaction terms (*p=0.01 and p=0.002, respectively*). We also analysed the variations in mean frame-wise displacement using principal component analysis (PCA) over runs. We found that the second principal component (PC-2) explaining 16% of the variation was associated to the alterations in mean frame-wise displacement during movie-watching condition. The projections of PC-2, related to movie-watching mean frame-wise displacement, were not correlated with the projections of condition-specific PC (*Spearman rank r=0.02, p=0.85*).

Apart from head motion, various other confounding factors may affect during movie-watching condition. For this reason, we repeated all the analyses after regressing out the global signal from the time-series of each ROI for each subject and run.

### Inter-subject synchronization

To establish the construct validity of the principal component topography, we compared the condition-specific PC with inter-subject synchronization functional connectivity (ISS-FC)(Kim et al., 2017; Simony et al., 2016). ISS-FC was proposed as a measure to remove the effects of spontaneous activity and to define inter-regional correlations based on common stimuli across subjects. To calculate ISS-FC, we randomly split the subjects into 2 groups (50 random groups) and calculated the average BOLD time-series of each region over subjects per group. Then, we calculated the correlations between the average BOLD time-series across pairs of regions. This procedure was repeated separately for 2 resting-state and 2 movie-watching runs, and the average ISS-FC across movie-runs were reported in the main results. Since the sample size in this study is small, we replicated the analyses in the previous studies (Kim et al., 2017) and demonstrated the ISS-FC at resting-state and movie-watching conditions (Supplementary Figure 2).

### Time-resolved functional connectivity

To extract time-resolved functional connectivity (dynamic FC; dFC), we used phase locking values (PLVs) of BOLD time series in a narrow frequency band (Demirtaş et al., 2016; Glerean et al., 2012). This approach enables the characterization of connectivity patterns at each time point, and it does not require specification of a window and a step-size, as in sliding-window analysis. The preprocessed time series were band-pass filtered in *0.04-0.07Hz* range to reduce the effects of low-frequency drift and high-frequency noise (Glerean et al., 2012). The Hilbert transform was then used to quantify instantaneous phase. The Hilbert transform, S(t) = Acos(φ(t)) of the preprocessed BOLD time series decomposes the signal into to an analytical signal *S(t)* with an instantaneous phase *φ(t)* and amplitude *A*. For each time point *t*, the difference Δ*φ*_*ij*_(*t*) between the phases of the respective ROIs was calculated, where *i* and *j* are the indices of each ROI. The phase differences were adjusted between 0 and *π* such that:

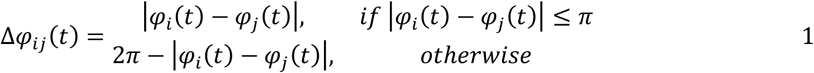

Then, the phase-locking values (PLVs), PLV_ij_(t) were constructed using the phase differences normalized between 0 and 1, thereby representing perfect anti-synchronization and perfect synchronization respectively, such that: PLV_*i,j*_(t) = 1 − Δ*φ*_*i,j*_(*t*)/*π*.

### PCA trajectories of time-resolved FC

The principal component analysis was repeated for grand average PLVs, to establish the link between Pearson correlation-derived FCs and PLVs. Since the PLVs were more sensitive to global synchronization levels, we subtracted the mean from each average PLV matrix before performing any analyses. After identifying the grand average condition-specific PLV component, we performed PCA on concatenated PLVs over time for each subject (i.e. 2 resting-state and 2 movie-watching runs). The condition-specific temporal components were identified as the PC with the highest similarity to the grand average condition-specific PLV component. We then characterized the trajectories (i.e. PC scores over time) of the condition-specific temporal components of the subjects. Here, the term “trajectory” was preferred over “scores” to highlight the fact that the PCA was performed over time. We asked whether the PC showing highest similarity to the condition-specific component distinguishes between resting-state and movie-watching trajectories. We quantified the condition-specific distinction by calculating the average distance between the median trajectories of the resting-state and movie-watching conditions. The distances between median trajectories were defined as the squared difference between median PC scores of resting-state and movie-watching trajectories. For each subject, we assessed the significance of the distinction by comparing the condition-specific distance against the surrogates. The trajectories of each subject were randomly shuffled and then assigned into two groups. The p-values were based on the distance between condition-specific trajectories, relative to the surrogate distances. Since the individual PC trajectories are time-dependent, we assessed the difference between conditions across subjects by calculating the median distances across conditions and runs. For each subject, the median trajectory distance between resting-state and movie-watching conditions was calculated. Then, the distances between 2 separate runs of resting-state and movie-watching conditions were calculated. Finally, we performed a permutation t-test to compare the average distance across conditions and runs.

### Time-resolved FC similarity across conditions and runs

To study the role of time-locked events on PLV dynamics during movie-watching condition (analogous to inter-subject synchronization), we calculated the similarity between instantaneous PLVs across conditions and runs. For each time point, we calculated the similarity between the PLVs of a single subject (k) and the average PLVs across the rest of the subjects (n≠k). The average PLVs were calculated to test the PLV similarity in 3 different circumstances: Across conditions (i.e. if subject k is at resting state run 1, the average PLVs were calculated for movie-watching run 1), across runs (i.e. if subject k is at resting state run 1, the average PLVs were calculated for resting state run 2) and within runs (i.e. if subject k is at resting state run 1, the average PLVs were calculated for resting state run 1).

### Computational modelling

We modelled the whole-brain rs-fMRI BOLD signals using 66 nodes. Each node was coupled with each other via DWI-derived structural connectivity (SC) matrix. We described the local dynamics of each individual node using normal form of a supercritical Hopf bifurcation (Deco et al., 2017). The advantage of this model is that it allows transitions between asynchronous noise activity and oscillations. Where ω is the intrinsic frequency of each node, a is the local bifurcation parameter, η is additive Gaussian noise with standard deviation β, the temporal evolution of the activity, z, in node j is given in complex domain as:

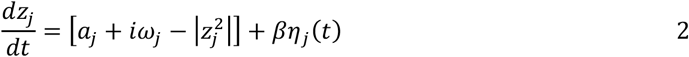

and,

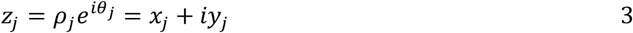

This system shows a supercritical bifurcation at *a*_*j*_ = *0*. Being specific, if *a*_*j*_ is smaller than 0, the local dynamics has a stable fixed point at *z*_*j*_ = *0*, and for *a*_*j*_ values larger than 0, there exists a stable limit cycle oscillation with a frequency *f = ω/2π*. Finally, the whole-brain dynamics is described by the following coupled equations:

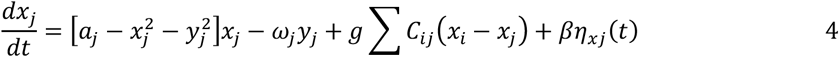

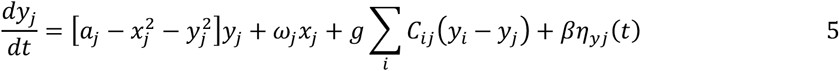

Where *C*_*ij*_ is the Structural Connectivity (SC) between nodes i and j, g is the global coupling factor, and the standard deviation of Gaussian noise, *β = 0.02*. The natural frequency (*f*) of each region was taken as the peak frequency in the given narrowband of the corresponding region in the empirical time-series.

Following a similar approach previously employed (Deco et al., 2014), we analytically estimated the model FC using linearization of the system around a stable fix point. Where *δ***u** = {*δx*_1_ …*δx*_66_, *δy*_1_ …*δy*_66_} represents the Taylor expansion of the system, **A** is the Jacobian matrix, and *ɛ*(*t*) is the noise term, the fluctuations around the fix point can be described as:

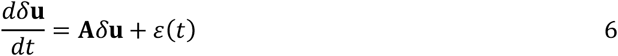

Where the deterministic parts of right-hand side of equations 4 and 5 are described by −*F*_*j*_ and −*G*_*j*_, respectively, the Jacobian matrix of the system evaluated at the fixed point 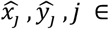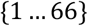 can be constructed as:

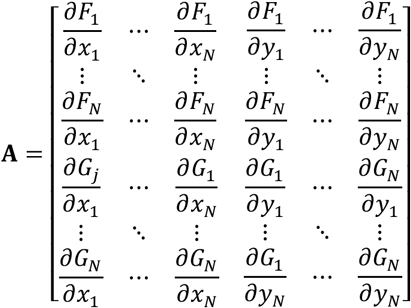

Where *i,j* ε {1 … 66}, each element of matrix **A** can be calculated as:

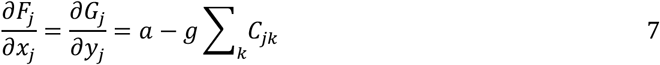

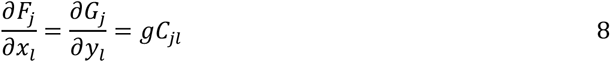

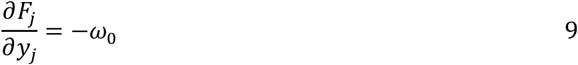

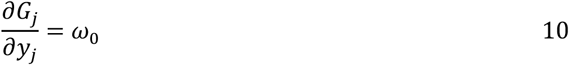

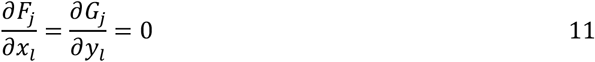

Where **Q** is the noise covariance matrix, the covariance matrix of the system **P** can be estimated by solving Lyapunov equation:

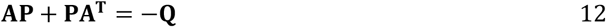

Finally, the model correlation matrix (FC) can be extracted from the covariance matrix as:

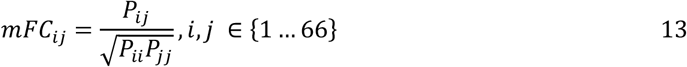

We estimated the model optimal parameters a and g by maximizing the similarity between model FC (equation 13) and empirical FC using gradient descent optimization. For each subject, the empirical functional connectivity was calculated as the average FC across the corresponding conditions (i.e. resting state or movie sessions) of the corresponding subject. The similarity between model FC and empirical FC was quantified as Pearson correlation similarity. To avoid the solutions reflecting a local minimum, for each subject/condition we estimated the best solution after repeating the optimization with 100 random initial conditions.

### Statistical analyses

The comparisons across conditions (resting-state versus movie sessions) were done using permutation t-test. Since the same subjects were tested under different conditions, we used dependent t-test. The randomization during the permutation t-test was also controlled to preserve this dependence across conditions. For optimal bifurcation parameters, the p-values were FDR corrected (*p<0.01*), with the Benjamini & Hochberg algorithm, when appropriate (Hochberg and Benjamini, 1990).

To assess the association between measures, we used Spearman rank correlations (to avoid potential contribution of outliers and due to limited sample size). Calculating the correlations separately for each condition (due to repeated-measures) did not alter the significance; therefore, for simplicity we reported a single correlation value between each measure. We used Pearson correlation as a measure of similarity between connectivity matrices (i.e. PC axes, FCs, PLVs).

The visualizations of the cortical surface were done using Connectome Workbench software. We used a population-average cortical surface (Conte69) (Van Essen, 2005), and a template to visualize the anatomical parcellations on the cortical surface.

## Acknowledgements

We thank Alan Anticevic, John Murray, Markus Helmer and Joshua Burt for the insightful discussions and their comments during the preparation of this paper.

## Notes

**Funding:** GD is supported by the Spanish Research Project PSI2016-75688-P (AEI/FEDER) and by the European Union’s Horizon 2020 Framework Programme for Research and Innovation under the Specific Grant Agreement. No. 785907 (Human Brain Project SGA2).

